# Quantification of differential toxin expressions and their relation to distinct lifespans of bacterial subpopulations associated with diverse host immune mechanisms

**DOI:** 10.1101/2022.10.11.511682

**Authors:** Shweta Santra, Indrani Nayak, Dibyendu Das, Anirban Banerjee

## Abstract

An assortment of robust intracellular defence mechanisms are critical for restricting proliferation of pathogens and maintaining sanctity of the cytosol. Defect in these mechanisms could be exploited by the pathogens for creation of a safe sanctuary which can act as a transient reservoir for periodic dissemination into the host. While residing inside the host cell, pore forming toxins secreted by the pathogens compromises the integrity of the vacuole and exposes the microbe to diverse intracellular defence mechanisms. However, the correlation between toxin expression levels and consequent pore dynamics, fostering pathogen’s intracellular life, remains largely unexplored. In this study, using *Streptococcus pneumoniae* (SPN) and its secreted pore forming toxin pneumolysin (Ply), as model systems, we explored various facets of host-pathogen interactions in host cytosol, governed by the toxin expression and the resultant pore formation. The extent of damage on the endosomal membrane was found to dictate subsequent interaction with different host endosomal damage sensors. This in turn governed the routes of SPN clearance, revealing multiple layers of defence mechanisms at host’s disposal for counteracting invaded pathogens. A subset of SPN population producing extremely low amount of Ply inflicted minimal damage to the endomembrane, precluding decoration by endomembrane damage sensors and significantly prolonging its intracellular persistence. Such long persisting bacterial population could be key for pathogenic transmission or ensuing invasive disease. Using time-lapse fluorescence imaging, we monitored lifespans of different pneumococcal population subsets inside host cells. After quantitative analysis of various timescales such as pore formation time, vacuolar or cytosolic residence time and total degradation time, we developed a mathematical model that could correlate these to intravacuolar accumulation of Ply monomers. By proposing events like pore formation and vacuolar degradation of SPN as first passage processes, our theoretical modelling yields estimates of Ply production rate, burst size, and threshold Ply quantities which triggers these outcomes. Collectively, we present a general method by which intracellular lifespans of pathogens could be correlated to differential levels of toxins that they produce.

## Introduction

Host intracellular defence mechanisms ensure effective pathogen elimination and maintain cytosolic sterility. However, many intracellular bacterial pathogens reside within a customized vacuolar compartment which not only allow bacterial replication but also protect the microorganism from cytosolic antimicrobial strategies [1, 2]. For quite a few pathogens, residence inside these vacuolar compartments is compromised due to expression of pore forming toxins by the pathogen. The loss of integrity of the vacuole exposes the microbe to diverse fail-safe intracellular defence mechanisms. Maintenance of the integrity of these vacuoles is therefore critical for prolonged persistence of these pathogens even though they may express pore forming toxins which could be essential for other aspects of their lifecycle [3]. But the dynamics of toxin expression and subsequent pore formation, promoting interaction with cytosolic immune systems leading to its clearance or evasion from such defences ensuring its safety and promoting intracellular life, is not known.

*Streptococcus pneumoniae* (SPN), the Gram-positive opportunistic pathogen, commonly resides in the upper respiratory tract of healthy individuals as commensal. But time to time it disperses from its niche and causes life-threatening diseases such as pneumonia, septicemia, endocarditis and meningitis [4]. Although typically considered an extracellular pathogen, it does have a brief intracellular stint which is particularly pertinent during trafficking across host barriers, such as the blood-brain barrier (BBB) and lung epithelial barrier [5] which are key aspects of its disease pathogenesis. In recent times, few studies have also identified that SPN replicates and persists for prolonged period inside splenic macrophages and this serves as a transient reservoir for dissemination into the blood causing septicemia [6]. Additionally, SPN has been shown to replicate within cardiomyocytes [7] and few serotype I SPN strains have been shown to adapt a benign intracellular niche in lung epithelium [8]. It remains elusive though whether these intracellular niches constitute a reservoir for SPN to re-establish colonisation and thereby facilitating transmission to new hosts. But these evidences have prompted the speculation that pneumococci may be evolving down an alternative evolutionary trajectory by embracing an intracellular lifestyle and induction of a milder form of disease for efficient transmission [8].

Central to this intracellular life of SPN is pneumolysin (Ply), the cholesterol dependent pore-forming toxin secreted by all serotypes of SPN, which promotes cell lysis and extensive tissue damage in the host [9]. Although Ply is reported to bind to mannose receptor C type 1 (MRC-1) and sialyl LewisX antigen [9], it’s cytolytic and inflammatory activity is primarily governed by membrane cholesterol binding. Like any other Cholesterol dependant cytolysin (CDC), 40 to 50 Ply monomers assemble on cell membrane to form a pre-pore complex which converts into ring-shaped functional pore of 200-500Å, sufficient enough to cause haemolysis [10–12]. But the pore sizes are not always of similar pattern or of equal diameter. High-speed AFM analysis has further revealed that diverse shapes such as arcs or slits can also form functional pores and still exhibit cytolytic activity [13]. Functional significance of such structural diversification among pore sizes is yet to be discovered. Moreover, these *in vitro* experiments performed with purified Ply does not provide the real scenario prevailing during SPN’s interaction with host where intravacuolar accumulation of Ply presumably governs the fate of the pathogen.

We have earlier established the key role of Ply in mediating interaction of SPN with intracellular defence mechanisms [14]. We demonstrated that Ply inflicted damage on endomembrane triggered recruitment of various ‘eat me’ signals like galectins or ubiquitin which then shunt SPN towards autophagy or ubiquitin mediated degradation pathways [15]. However, correlation between expression levels of Ply by SPN in the intravacuolar environment governing pore dynamics and subsequent choice of fates fostering its intracellular life was not established. In this study, we quantitatively relate the dynamics of pore formation and SPN degradation to a model of Ply gene expression which provides an estimate of intravacuolar Ply accumulation dictating its fate. A threshold crossing of Ply quantity is a first passage event which in general refers to the occurrence of a relevant event for the first time in a stochastic process. Several cellular events in bacteria and yeast have been modelled in recent years as first passage processes. For example, lysis of *E. coli* cells due to the accumulation of λ-phage Holin protein beyond threshold quantity has been mathematically treated as a first passage process, and its time statistics have been extensively studied [16–21]. Various other biological events such as, the timing of kinetochore capture by microtubules of a fission yeast, accumulation of a critical cell division protein FtsZ during cell division in *E. coli* has also been treated as the first passage [22–24]. However, no such quantitative studies have been performed on any intracellular pathogen’s degradation time using this approach. Since Ply expression is established to be stochastic, it’s accumulation on endomembrane beyond a threshold, forming variety of pores, can be considered as a first passage process. Here, using stochastic modelling, we provide quantitative estimate of the differential rate of Ply expressions in two distinct subpopulations of SPN, suffering degradation by two distinct pathways, e.g., vacuolar and cytosolic. Our analysis not only revealed the contribution of Ply expression regulating the bacterial life spans inside the host, but also revealed the strategy that SPN adopts to secure a safe intracellular life for prolonged periods.

## Materials and Methods

### Bacterial strains

SPN strain R6 (Serotype 2, gift from Prof. T J Mitchell, University of Birmingham, UK, Retd.) and its derivatives were routinely grown in Todd-Hewitt broth (THB) supplemented with 1.5% yeast extract at 37□ in 5% CO_2_. When necessary following antibiotics were used: Spectinomycin (100 μg/ml), Chloramphenicol (4.5 μg/ml). Generation of engineered pneumococcal strains, such as, Δ*ply* mutant, low (SPN:Ply-Low) and high (SPN:Ply-High) Ply expressing strains, have been described earlier [15].

SPN strains were made fluorescent either by integration of *hlpA*-tRFP fusion cassette (Kind gift from Prof. JW Veening, University of Groningen, Netherlands) into the genome of SPN or by staining with DRAQ5 (BD Biosciences) and Hoechst (HiMedia). All gene deletions and cassette insertions were verified by PCR amplification of the gene locus, followed by DNA sequencing.

### Cell culture and transfection

Human lung alveolar adenocarcinoma (Type II pneumocyte) cells or A549 (ATCC No. CCL-185) were routinely cultured in DMEM medium (HiMedia) supplemented with 10% Fetal Bovine Serum (FBS, Gibco) at 37□ and 5% CO_2_. Stably transfected A549 cells overexpressing YFP-Gal8, GFP-LC3, mStrawberry-Galectin3, mCherry-LyseninW20A fusion proteins were generated following transfection with M6P-Blast-YFP-LGALS8 (kindly provided by Prof. Felix Randow, MRC Laboratory of Molecular Biology, UK), pMRX-IRE-Blast-GFP-LC3 (subcloned from pBABE-Puro-GFP-LC3 vector, Addgene # 22405), pMRX-IRE-Puro-mStrawberry-Galectin3 (kindly provided by Prof. Tamotsu Yoshimori, Osaka University, Japan) and pMRX-IRE-Blast-mCherry-LyseninW20A (subcloned from M6P-blast-mcherry-lyseninW20A, kindly provided by Prof. Felix Randow, MRC Laboratory of Molecular Biology, UK). All transfections were performed using Lipofectamine 3000 (Invitrogen) as er manufacturer’s instructions. Following transfection, cells were selected with 2 μg/ml of puromycin (Sigma Aldrich) and 2 μg/ml of blasticidin hydrochloride (HiMedia). For treatment studies, cells were incubated in 50 μM of monensin sodium salt (Sigma Aldrich) post infection.

### Cell viability assay

To find out the working concentration of monensin, A549 cells were incubated with varied concentration of monensin from 50 μM to 300 μM. At 9 h post incubation, cell viability was checked using MTT assay kit (HiMedia). Cells incubated with ethanol (vehicle) or Triton X100 (0.1%) were considered as negative and positive controls, respectively.

### Gentamycin protection assay

Mid-log phase grown SPN strains were re-suspended in PBS to adjust the OD_600nm_ to 0.4 comprising of 10^8^ cfu/ml. These were used to infect A549 cells with MOI 25 and incubated for 1 h followed by treatment with penicillin (10 μg/ml) and gentamicin (400 μg/ml) to kill extracellular SPN. At indicated time points cells were thoroughly washed with DMEM, lysed with PBS containing 0.025% Triton X-100 and the lysates were plated on BHI agar plates to enumerate survived bacteria. Percentage survival at indicated time points were calculated relative to 0 h and survival efficiency at indicated time points was represented as fold change in percent survival relative to control.

### Fluorescence imaging

For immunofluorescence, A549 or its transfected derivatives grown on glass coverslips were infected with SPN at the multiplicity of infection (MOI) of 25 for 1 h followed by treatment with penicillin (2 μg/ml) and gentamicin (400 μg/ml) for 2 h to kill the extracellular bacteria. At desired time points, infected cells were washed several times with DMEM medium, fixed with 4% paraformaldehyde for 15 min, permeabilized with 0.1% Triton X-100 for 10 min and blocked with 3% Bovine serum albumin (BSA) for 2 h. Cells were incubated overnight in appropriate primary antibodies prepared in 1% BSA in PBS at 4□. Next day, cells were washed with PBS and incubated in suitable secondary antibodies for 1 h. Finally, cells were incubated in Hoechst (10 μg/ml) for 15 min and mounted over a clean glass slide using VectaShield® without DAPI (Vector Laboratories) for visualization using a Laser Scanning Confocal microscope (LSM 780, Carl Zeiss) under 40X or 63X oil objectives. Images were acquired after optical sectioning and processed using Zen Lite software (Version 5.0).

Live fluorescence imaging was performed in a Spinning Disk Confocal Microscope or Laser Scanning Confocal Microscope. Monolayer of transfected cells grown in a 35 mm glass-bottom petri dish were infected with DRAQ5 (3 μg/ml) treated SPN as mentioned above. Time-lapse imaging was performed at multi-position along with optical sectioning under 63x oil immersion objective and images acquired were processed and analyzed using Zen Lite Software 3.1.

### Measurement of pH

For generating of a pH calibration graph, cell monolayer grown in 35 mm glass-bottom dish was incubated in FITC (200 μg/ml) for 1 h at 37□ and 5% CO_2_. Post incubation, cells were extensively washed followed by sequential incubation in different isotonic K^+^ solutions (140 mM KCl, 1 mM MgCl2, 0.2 mM EGTA, 20 mM NaCl and 20 mM of either HEPES or MES or Tris based on pH) buffered to pH values from 4.5 to 7.5 containing 5 μM monensin. After 5 min, ratiometric imaging of cells was performed in Laser Scanning Confocal Microscope using 63X oil immersion objective. Light was transmitted alternately through 488 nm and 405 nm excitation filter while the emitted light was passed through 520 nm emission filter and captured using GaSP detector. The resulting fluorescence intensity ratio (490 / 440) was extrapolated to its corresponding pH values. For measuring pH of SPN containing endosomes, cells were infected with SPN in a FITC laden media and ratiometric imaging of bacteria possessing endosomes was performed as mentioned.

### Antibody and Inhibitors

The following antibodies were used in this study were Galectin3 (R&D systems, 194805), Galectin 8 (R&D systems, AF1305), anti-goat IgG 633 (Invitrogen, A21082), anti-mouse IgG 555 (Invitrogen, A21422). Fine chemicals used were DiO (Invitrogen), Monensin sodium salt (Merck), DRAQ5 (BD Biosciences), FITC (Sigma Aldrich), Bafilomycin A1 (Merck), Flipper-TR (Cytoskeleton Inc.).

### Theoretical formula for threshold crossing times of Ply number

We intend to relate the occurrence of the cytosolic escape of certain subpopulation of SPN through pore formation in Pneumococcal Containing Vacuoles (PCVs), as well as the degradation of other long-lived subpopulations inside PCV, to the level of Ply expressions and their crossing of thresholds. For Ply expression, we assume a model with the following features used for protein expressions in bacteria [25, 26];

- Proteins are produced in bursts and the burst-size follows a geometric distribution [25–28]. The Poisson rate (*r*) of making a transition from a state with protein number *i* to a state with protein number (*i* + *n*) is given by 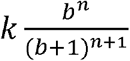. Here, *k* is the transcription rate assumed to be independent of the instantaneous protein number (*i*), i.e., there is no feedback based on protein number. The mean burst size is *b*, while the change in protein number *n* ∈ [0, 1, 2, …, ∞].
- The lifetimes of mRNA are typically much shorter than the lifetime of proteins. Within the timescales of the cellular events of our concern (namely SPN degradation), the proteins (Ply) are assumed not to degrade.
- Mathematically the time evolution of probability *P_n_* of the protein number *n* is given by the forward Master equation [25]:

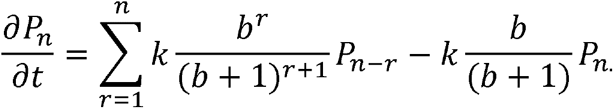

Our concern is the statistics of timescales for the Ply number *n* to start from an initial value *N_0_* and stochastically increase and cross a threshold value *N* for the first time. Recently this mathematically problem on first passage times has been analytically solved [20, 21] and the exact distribution of threshold crossing times *t* [20] is as follows:

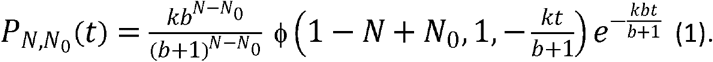

Here ϕ is a confluent hypergeometric function [29]. The theoretical mean and standard deviation of the threshold crossing times (from Eq. (1)) are

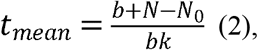

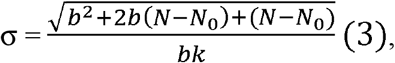

respectively [20].

### Procedure of plotting histograms of the experimental data and fitting with theoretical formulas

From live-cell images, we quantified time representing three kinds of events observed in the degradation pathways of different subpopulation of WT SPN. For the subpopulation which degrades in the cytosol, the relevant event timescales are as follows: (i) Gal8 attachment times *t_g8_*, which are seen for both cytosolic and vacuolar degradation pathways; (ii) time for formation of large pores on the Gal8 marked PCVs for cytosolic escape *t_p_* (applicable for cytosolic degradation pathway); (iii) lifetime of SPN within the autophagosome after Gal8 recruitment *t_v_* (in case of vacuolar pathway).

The Gal8 attachment depends on sensing a minimal rupture in the membrane of PCV caused by Ply. Since attachment process is expected to be a single step binding, the statistics of the timescales *t_g8_* is given by *P*(*t_g8_*) which follows exponential distribution. On the other hand, the cytosolic escape time *t_p_* is expected to depend on Ply accumulation on the inner membrane of autophagosome and its number crossing a certain threshold. Hence, *t_p_* is distributed according to the formula in Eq. (1). Likewise, for vacuolar degradation, the vacuolar degradation time *t_v_*, after Gal8 recruitment, is also expected to be distributed according to Eq. (1).

For the quantitative analysis of data, normalized histograms of the above time points were plotted using MATLAB software. We fit these histograms with normalized exponential distributions ~ λ exp (-λ *t_g8_*). The inverse of means of *t_g8_* (80 min and 137 min for the vacuolar and cytosolic cases, respectively) are used as the decay constants λ for the fitting.

For cytosolic degradation, we plotted the histograms for total cytosolic escape times, *t_C’_*=*t_g8_*+*t_p_*, and *t_p_*. Note that for every cytosolic escape event *t_C’_*>*t_g8_*, so *t_p_* = *t_C’_*-*t_g8_ >0*. We fit the histograms of *t_p_* and *t_C’_* using the theoretical formula mentioned in Eq. (1). We first fixed a burst size *b*. We then equated *t_mean_* in Eq. (2) with the experimental mean *t_p_* and varied (*N-N_0_*) as a free parameter (integer), the rate *k* gets fixed from Eq. (2). For our choice *b=2*, and certain (*N-N_0_*) and *k* as mentioned in the text, we found that the error between theoretical fit in Eq. (1) and experimental histogram in terms of SSR (sum of squared residuals at mid time points of each bin in the histograms) [30] is the least. We evaluated the σ from Eq. (3) for the best fitted values of *b, k* and (*N-N_0_*). For the vacuolar degradation pathway, we plotted histograms of total degradation times, *t_Vac_*= *t_g8_+ t_v_* and *t_v_*. Again note that for every vacuolar degradation event *t_Vac_* > *t_g8_*, so *t_v_= t_Vac_-t_g8_* > 0. To best fit the histogram of *t_v_*, fitting method similar to *t_p_* is used, as described above. To verify the goodness of the fit, the standard deviation σ from theoretical fit and experimental histogram were compared.

### Statistical analysis

GraphPad Prism version 5 was used for statistical analysis. Statistical tests undertaken for individual experiments are mentioned in the respective figure legends. p<0.05 was considered to be statistically significant. Data were tested for normality and to define the variance of each group tested. All multi-parameter analyses included corrections for multiple comparisons and data are presented as mean□±□standard deviation (SD) unless otherwise stated

## Results

### Heterogeneous pneumolysin expression drives differential degradation kinetics of SPN

An isogenic set of intracellular SPN gives rise to different intracellular subpopulations with respect to recruitment of various host-specific markers onto PCVs. This differential recruitment of various markers was attributed to heterogeneous expression of Ply [15]. However, the real-time kinetics of their recruitment and degradation dynamics of SPN in these PCVs remains unexplored. We therefore monitored the tRFP expressing WT SPN trapped inside YFP-Gal8 compartment by time-lapse confocal microscopy for extended hours post infection. Expectedly two distinct phenomena based on their degradative fate were evident in our experiments. In one phenomenon, the tRFP signal of SPN gradually faded inside the Gal8 vacuole, suggesting probable degradation of SPN inside Gal8 marked autophagosome (Fig. 1A-C). Gal8 is an endosome damage sensing marker which interacts with autophagy marker LC3, ensuring sequestration of the damaged endosomes in autophagosomes and subsequent degradation of the cargo following fusion with lysosomes [15, 31]. Quantitative analysis of a series of similar time-lapse imaging events (n = 50) revealed that while the mean Gal8 recruitment time (*t_g8_*) on PCVs was 80.4 ± 15.95 min, the average lifespan of SPN inside these Gal8 marked compartments (*t_v_*) was 184.84 ± 19.49 min (Fig. 1D-E). Combined together the mean lifetime (*t_Vac_*) of SPN in the Gal8 marked vacuolar environment was found to be 265.24 ± 21.39 min (Fig. 1F).

**Fig 1:**
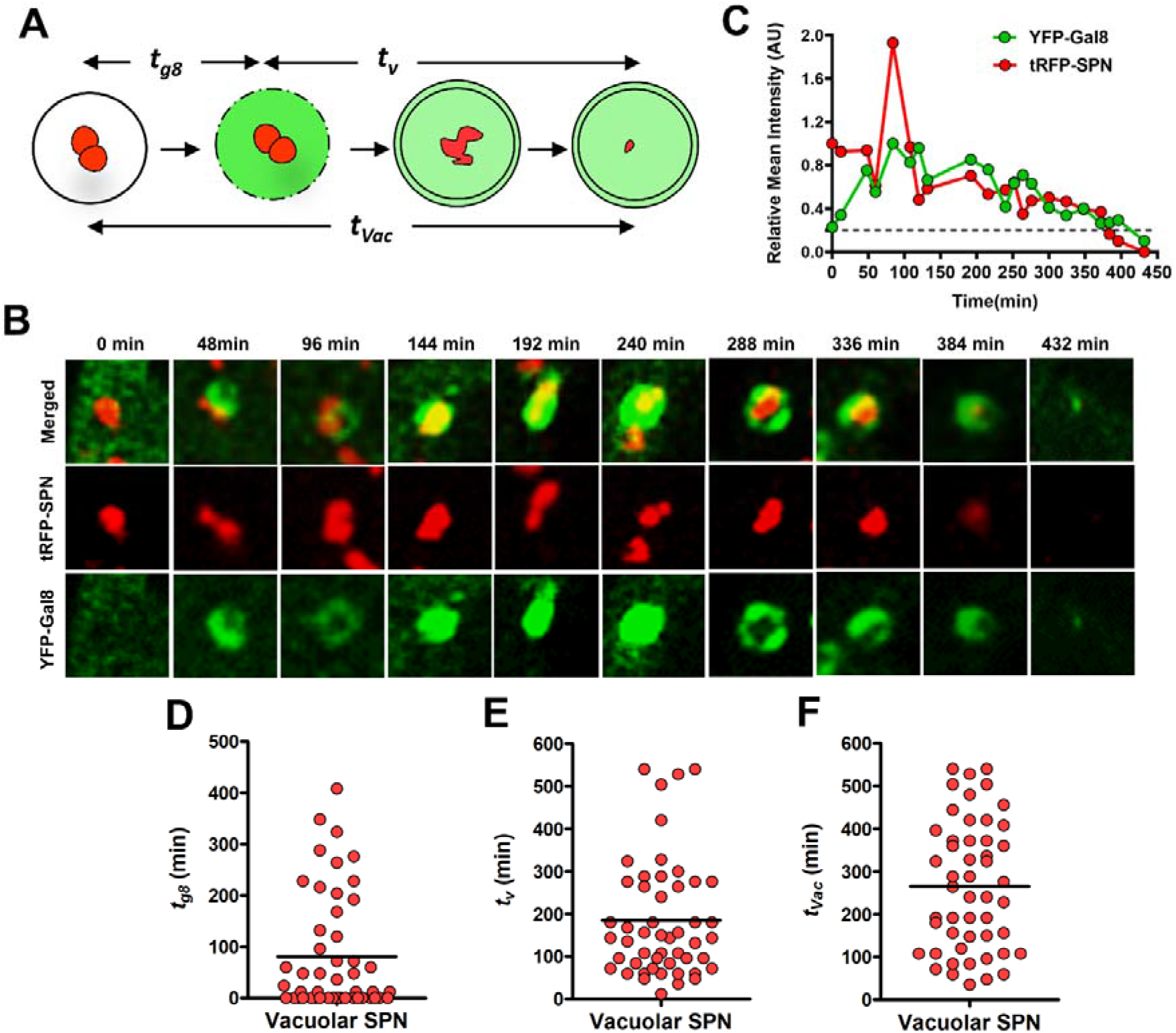
Temporal kinetics of vacuolar degradation of SPN. **(A)** schematic representation of SPN killing inside Gal8 marked compartment. *t_g8_*, Gal8 association time; *t_v_*, degradation time in Gal8 marked autophagosomes; *t_Vac_*, total degradation time for vacuole bound SPN. **(B)** Representative time-lapse montage depicting SPN degradation inside Gal8 marked compartments. A549 cells stably expressing YFP-Gal8 (green) were infected with WT SPN constitutively expressing tRFP (red) and live imaging of infected cells was performed with intervals of 12 min for extended hours using a confocal microscope. The stills correspond to the movie shown as S1 Movie. **(C)** Comparison of fluorescent intensities between Gal8 (green) and SPN (red) relative to its initial fluorescence intensity (*I_t’_/I_t0_*, where *t′* and *t_0_* represent the intensities of both signals at time 0 min and the given time point, respectively). **(D-F)** Scatter plot diagrams depicting the time of Gal8 recruitment on PCVs (**D**), time taken for degradation of SPN in Gal8 marked compartments (**E**) and total time of their residence in a vacuole (**F**), respectively.

While 55.5% of SPN were eliminated inside Gal8 marked autophagosomes, for a significantly high proportion (44%) of pneumococci there was catastrophic damage of Gal8 structures, triggering escape of SPN into the cytosol (Fig. 2A-C). Such sudden disappearance of Gal8 marked compartments could be attributed to its extensive endomembrane damage due to high Ply expression. Kinetic measurements of large number of such events (n = 40), we noticed that in these populations, average time of Gal8 recruitment (*t_g8_*) was 137.4 ± 19.35 min, and within 69.9 ± 7.072 min of Gal8 recruitment, the vacuoles collapsed (*t_p_*), allowing SPN to become cytosolic. In cytosol, these pneumococci were ubiquitinated and after persisting for 75.85 ± 10.64 min in the cytosolic milieu (*t_c_*) (Fig. 2D-F), they were finally degraded either via autophagy or proteasomal pathway depending on the type ubiquitin chain topologies formed [32]. Combining all these, the average lifetime (*t_Cyt_*) of these SPN population was 283.15 ± 18.21 min (Fig. 2G).

**Fig 2:**
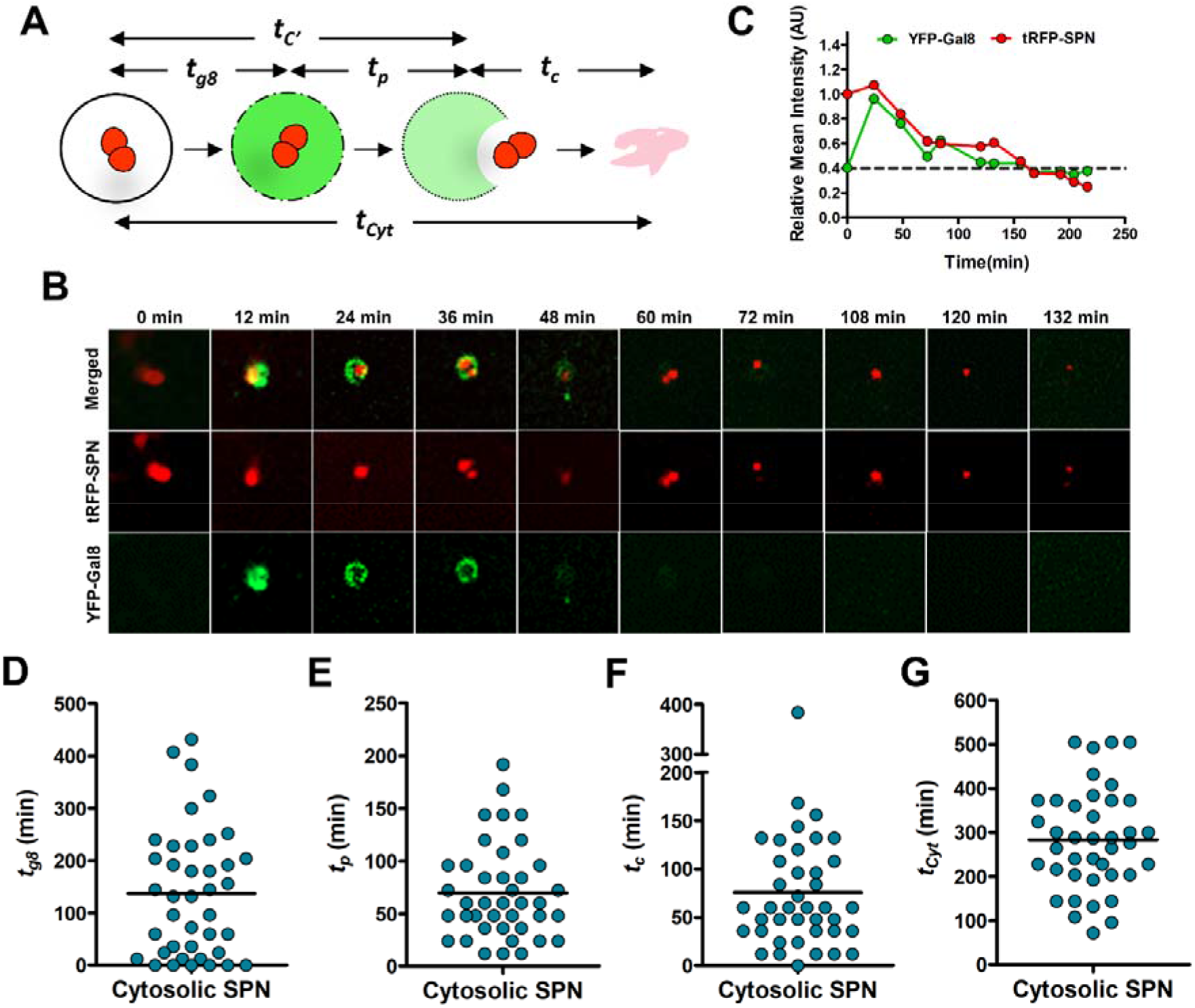
Kinetics of cytosolic SPN degradation in cytosol. **(A)** Schematic representation of SPN killing in cytosol. *t_g8_*, Gal8 association time; *t_p_*, time required for PCV puncture and release of SPN in cytosol; *t_c,,_* degradation time for SPN in cytosol; *t_Cyt_*, total time required for cytosolic bound SPN elimination. **(B)** Representative time-lapse montage of SPN escaping from Gal8 compartment and being degraded gradually in cytosol. A549 cells stably expressing YFP-Gal8 (green) were infected with WT SPN constitutively expressing tRFP (red) and time-lapse imaging of infected cells was performed with intervals of 12 min for extended hours under a confocal microscope. The stills correspond to the movie shown as S2 Movie. **(C)** Comparison of fluorescent intensities between Gal8 (green) and SPN (red) relative to its initial fluorescence intensity (*I_t′_/I_t0_*, where *t′* and *t_0_* represent the intensities of both signals at time 0 min and the given time-point, respectively). **(D-G)** Scatter plot diagrams depicting the time of Gal8 recruitment on PCVs (**D**), time of catastrophic rupture of vacuole leading to SPN’s escape to cytosol (**E**), time of residence of SPN in cytosol (**F**) and total time of cytosol bound SPN degradation (G).

### High Ply expression triggers pH imbalance

While closely analyzing the clearance kinetics of different SPN populations, we noticed a discrepancy in Gal8 recruitment timings between vacuolar and cytosolic population subsets. The average time of Gal8 association (*t_g8_*) for vacuolar events (80.4 ± 15.95 min) was ~ 1.7 fold quicker than those of cytosolic events (137.4 ± 19.35 min). This was dubious as eventual cytosolic population presumably would express higher Ply, triggering quicker pore formation and thus should have faster Gal8 association. To further shed light on this paradoxical observation, we first created two genetically modified strains of SPN expressing either high (SPN:Ply-High) or low (SPN:Ply-Low) amount of Ply [33] and infected A549 cells with these two SPN strains. Expectedly, SPN: Ply-High strain damages the endomembrane and escape into the cytosol, while SPN:Ply-Low strain is predominantly present in PCVs (Suppl. Fig. 1A). However, at initial stages post endocytosis, much before the collapse of the endomembrane leading to SPN’s cytosolic escape, we speculated that quick small pores (Gal8^-^) formed by SPN:Ply-High strain could create pH imbalance inside PCVs, temporarily delaying its acidification. But such minutely damaged PCVs may eventually be acidified by the action of vATPase, which is critical for further activation and larger pore formation by Ply. We hypothesized that this delay in acidification of SPN:Ply-High containing PCVs and subsequent activation of Ply to form larger pores capable of recruiting Gal8, required longer time (Fig. 3A). To prove this, we selected Lysenin as a marker for very minute pores which is reported to result in transbilayer movement of sphingolipids (Lys^+^) without exposing the luminal glycans for decoration with Gal8 (Gal8^-^) [34]. Irrespective of Ply expression status, at early time points post infection, we observed presence of PCVs with minute functional pores (Lys^+^Gal8^-^) as well as non-functional pores (Lys^-^Gal8^-^) (Fig. 3B-C). Surprisingly, just after invasion, a significantly high proportion (66.4%) of vacuole bound high Ply expressing SPN (SPN:Ply-High) was found to be Lys^+^Gal8^-^ which is almost 1.5 folds higher than that of SPN:Ply-Low (Fig. 3D). Contrarily, 55.4% of PCVs resulting from infection with SPN:Ply-Low was observed to be Lys^-^Gal8^-^, which is almost 2.08 fold higher than similar population exhibited by SPN:Ply-High (Fig. 3E). This implied that at early time points post infection, low Ply expression does not lead to formation of any pores, whereas SPN:Ply-High though can efficiently generate functional pores on endosomes, but such pores are small enough to not expose the luminal galactosides for their successful recognition by Gal8. We next measured pH of PCVs containing SPN:Ply-High or SPN:Ply-Low, that are Lys^+^Gal8^-^ and Lys^-^Gal8^-^, respectively. Expectedly, the average pH of the Lys^+^Gal8^-^ population was found to be near neutral (6.5) while that of Lys^-^Gal8^-^ was acidic (5.3) (Fig. 3F). Collectively, these suggest that quicker pore formation by high Ply expression induces pH imbalance which may have rendered rest of the Ply inactive. Upon restoration of acidic pH at a later time by vATPase, the process of membrane perturbation resumes, triggering delayed decoration by Gal8. To prove the role of vATPase mediated restoration of acidic pH in the PCV resulting in activation of Ply and formation of larger pores (depicted by getting marked with Gal8), we treated the cells with Bafilomycin A1, an inhibitor of vATPase. We observed 40% reduction of Gal8 association with PCVs formed by SPN:Ply-High (Fig. 3G). This proved the necessity of vATPase mediated lowering of vacuolar pH for Ply activation and larger pore formation triggering Gal8 recruitment.

**Fig 3:**
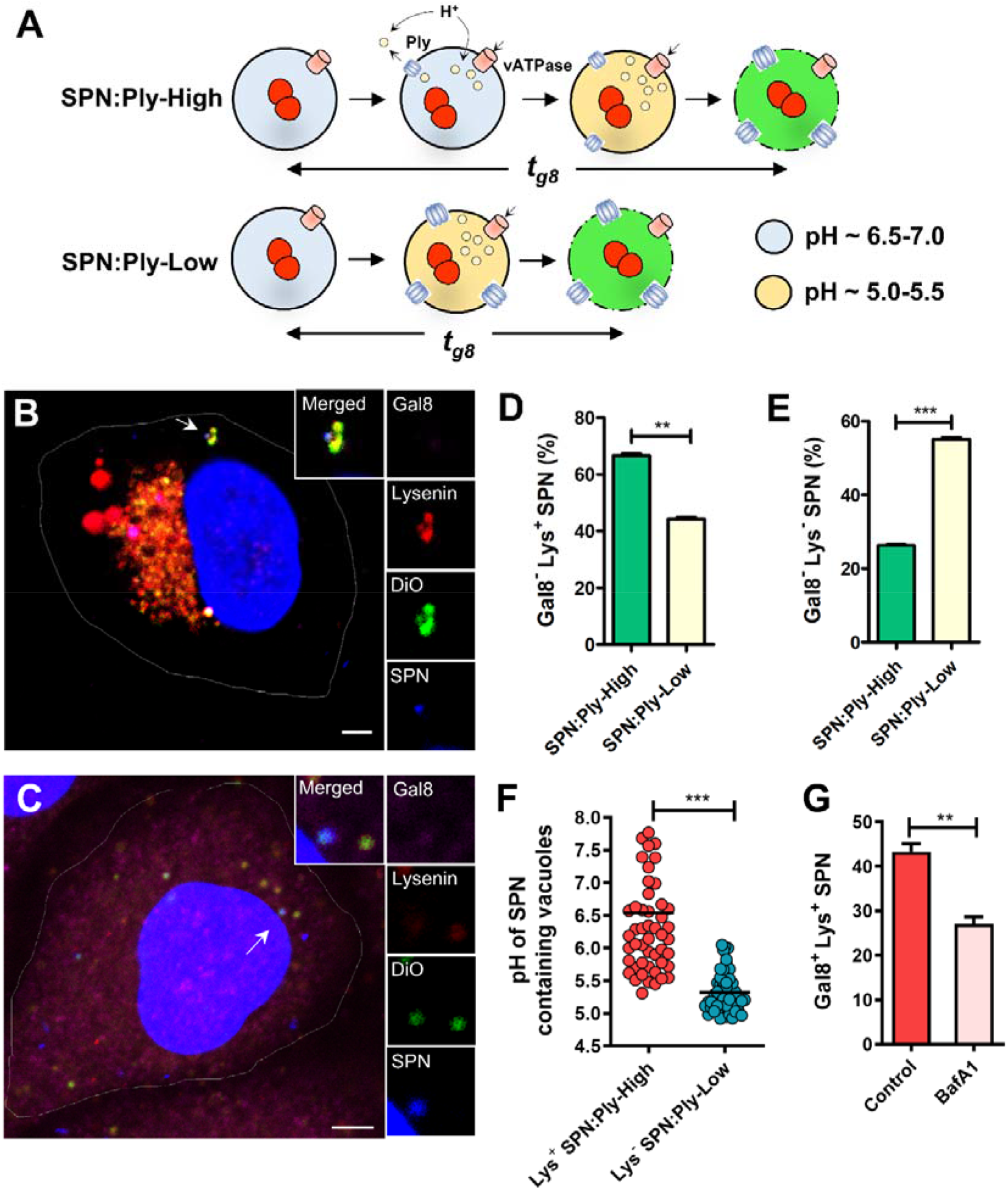
High pneumolysin expression produces quicker functional pores causing pH imbalance. **(A)** Schematic representation of Ply amount and pore dynamics. Following endocytosis, initial small pores formed by SPN:Ply-High strain temporarily delays endosomal acidification. Such minutely damaged PCVs may eventually be acidified by the action of vATPase, which is critical for activation and larger pore formation by Ply, capable of recruiting Gal8. SPN:Ply-Low on the other hand forms no pores to begin with due to less Ply, allowing acidification of PCVs and accumulation of Ply which suddenly may get activated to result in large enough pores for Gal8 recruitment. **(B-C)** Confocal micrographs showing association of SPN (blue) with lysenin (red), Gal8 (pink) and DiO (green) at 2 h post infection, DiO^+^Lys^+^Gal8^-^SPN (**B**) and DiO^+^Lys^-^Gal8^-^SPN (**C**). A549s stably expressing mChrerry-Lysenin was infected with SPN and stained with anti-Gal8 Ab, DiO and Hoechst. Arrowhead designates the bacteria shown in insets. Scale bar, 5 μm. **(D-E)** Quantification of co-localization of Gal8 negative PCVs containing SPN:Ply-High or SPN:Ply-Low with Lysenin at 2 h post infection. n ≥ 100 bacteria per coverslip. Data are presented as mean ± SD of triplicate experiments. **(F)** pH of PCVs at 2 h containing SPN:Ply-High or SPN:Ply-Low strains that are marked with or without Lysenin in A549 cells based on ratiometric fluorescence microscopy (n = 50 for each population). Fluorescence intensity ratios were converted to pH-values by fitting of data to pH calibration curve. **(G)** Percentage of decoration of PCVs (Lys^+^) containing SPN:Ply-High with Gal8 (Gal8^+^Lys^+^) following treatment of cells with Bafilomycin A1 (150 nM, 2 h). n ≥ 100 bacteria per coverslip. Data are presented as mean ± SD of triplicate experiments. Statistical analysis was performed using Students t-test (**D, E, F, G**). \*\**p*<0.005, \*\*\**p*<0.001.

### Less Ply expression facilitates prolonged intracellular persistence

Ply mediated larger pore formation is proven to be detrimental for SPN as following cytosolic exposure it is efficiently eliminated by host cytosolic surveillance and bacterial clearance mechanisms [15]. We therefore presumed that remaining confined within PCVs could be a preferred option for prolonged intracellular persistence. Such lifestyle is possibly ensured by less Ply expression. Indeed, SPN:Ply-Low exhibited ~ 4 fold higher survival ability compared to SPN:Ply-High strain (Suppl. Fig. 1B). Critically, 75% of low Ply expressing SPN were confined within vacuoles that were marked by autophagy marker LC3, but were devoid of endomembrane damage sensing marker Gal8 (Gal8^-^LC3^+^) (Suppl. Fig. 1C).

A recent report suggested Galectin 3 (Gal3) also to be an endomembrane damage sensor where it’s role was found to be pivotal for shunting the damaged endosome either towards repair pathway or sequestration by autophagosome [35, 36]. This suggested that the LC3^+^Gal8^-^ SPN population we observed for SPN:Ply-Low strain, which presumably have minute damage in the endomembrane, could be marked with Gal3. Indeed, we observed that majority (~10%) of PCVs containing SPN:Ply-Low were marked with only Gal3 and did not acquire Gal8 (Gal3^+^Gal8^-^) (Suppl. Fig. 2A, B). On the contrary, 37% of SPN:Ply-High harbouring PCVs were dually decorated (Gal3^+^Gal8^+^) (Suppl. Fig. 2A, C). This implies that while initial minute damage to endomembrane triggers Gal3 recruitment, subsequent major damage leads to Gal8 association.

Since majority of SPN:Ply-Low population are primarily decorated with Gal3, instead of Gal8 (due to formation of small pores), we then evaluated the role of Gal3 in driving SPN:Ply-Low strain towards autophagic killing. Surprisingly, we noticed that intracellular SPN:Ply-Low gave rise to three distinct subpopulations with respect to the markers associated with it (Suppl. Fig. 3A-C). While Gal3^+^LC3^-^ implies that this population subset is driven towards endomembrane repair pathway, the Gal3^+^LC3^+^ depicts targeting of SPN subset towards Gal3 mediated autophagic clearance. Although we observed both Gal3^+^LC3^-^ (6%) and Gal3^+^LC3^+^ (9%) marked vacuoles, a significant subset (23%) of SPN:Ply-Low were found to be decorated with LC3 but were devoid of even Gal3 (Gal3^-^LC3^+^) (Suppl. Fig. 3D). This underscores that recruitment of LC3 in this SPN population is not dependent on endomembrane damage sensing, but could possibly be attributed to pH/osmotic imbalance created by extremely small pores produced by SPN:Ply-Low strain, as has been suggested for VacA toxin producing *H. pylori* [37].

We therefore explored the fate of these different SPN populations in A549 cells stably expressing mStrawberry-Gal3 and GFP-LC3. For both Gal3^+^LC3^-^ and Gal3^+^LC3^+^ population subsets, we observed eventual fading of SPN signal, confirming degradation of the pathogen (Fig. 4A-D). The mean survival time for these two population subsets were 199 and 267 min, respectively (Fig. 4G). Critically, the Gal3^-^LC3^+^ population subset which represent the largest fraction of the SPN:Ply-Low population, survived till 9 h and beyond (Fig. 4E-G). Throughout the course of time lapse imaging, these SPN population subset remained associated with LC3 and were never marked with Gal3.

**Fig 4:**
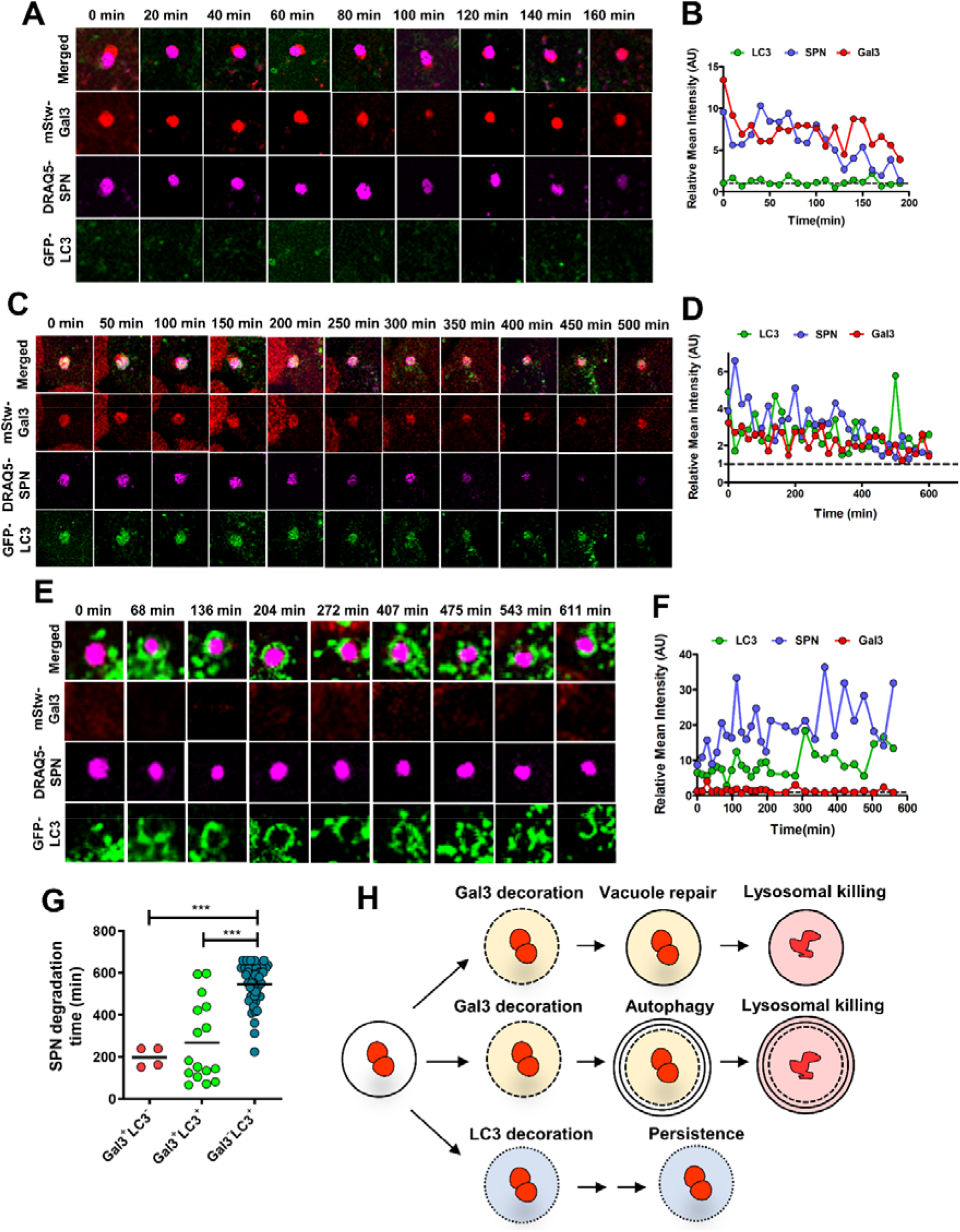
Differential fate of low ply expressing SPN. **(A), (C)** and **(E)** depict representative time-lapse montages of lifetimes of SPN inside Gal3 **(A)** or LC3 **(E)** or dually **(B)** marked vacuoles. A549s stably expressing mStrawberry-Gal3 and GFP-LC3 were infected with DRAQ5 stained SPN and time-lapse imaging was performed at 30 min post infection. The stills in A, C and E correspond to the movies shown as S3 Movie, S4 Movie and S4 Movie, respectively. **(B**, **D** and **F)** Temporal quantification of Gal3, LC3 and SPN fluorescence intensities associated with PCVs either relative to fluorescence intensity of the cytosol (*I_PCV_/I_Cyt_*) (for LC3 and Gal3) or fluorescence signal of host cell nuclei (*I_SPN_/I_Nuc_*) (for SPN). Relative mean intensity values ~ 1.0 for any marker means disappearance of the signal of the corresponding marker. In **(A)** and **(C)** relative mean intensity values for bacteria (*I_SPN_/I_Nuc_*) remains > 1.0 for long period of time before finally disappearing. However, in **(E)** *I_SPN_/I_Nuc_* remained > 1.0 throughout the course of imaging, highlighting prolonged persistence of low Ply producing SPN in only LC3 marked compartments. **(G)** Mean degradation time of SPN:Ply-Low population subsets as determined by the time lapse fluorescence imaging. Statistical analysis was performed using one-way ANOVA followed by Tukey’s Multiple Comparison Test. \*\*\**p*<0.001. **(H)** Schematic representation of different SPN population subsets arising due to expression of low Ply. In one population subset, following minute damage, Gal3 recruitment triggers endomembrane repair pathway which eventually results in lysosomal fusion and SPN killing. In another subset, damage followed by Gal3 association triggers autophagic sequestration of damaged PCV and subsequent lysosomal killing. Finally, extremely minute damage in the endomembrane leads to LC3 lipidation without association of any damage sensing markers and this population exhibits prolonged persistence.

### Induction of osmotic imbalance is sufficient for LC3 lipidation of PCVs and prolonging intracellular persistence

We hypothesized that the Gal3^-^LC3^+^ population have originated due to extremely low Ply synthesis resulting in negligible damage in the endomembrane. Such small pores are not sufficient for recognition by damage sensors like Gal3 but are adequate to create pH imbalance, promoting non-canonical LC3 lipidation of PCVs and simultaneously halt PCV maturation to prolong SPN’s persistence. To prove this, we infected A549 cells with a Δ*ply* mutant SPN strain, which does not possess any pore forming ability and therefore did not associate with LC3 or any other damage sensing markers, such as Gal3 or Gal8 (Fig. 5A). To mimic low ply mediated osmotic imbalance in the PCVs containing Δ*ply* mutant strain, A549 cells were treated with monensin sodium salt (50 μM), a Na^+^/H^+^ ionophore after infection with Δ*ply* mutant strain. We observed that with increasing non-cytotoxic conc. of monensin (Fig. 5C), higher percentages (37%) of Δ*ply* containing PCVs were marked with LC3, but they were devoid of damage sensor Gal3 (Fig. 5B). Further, time-lapse imaging showed that these SPN (LC3^+^Gal3^-^ Δ*ply*) continued to persist till 10 h and beyond (Fig. 5D-E). These unambiguously prove that occurrence of osmotic imbalance either artificially by using an ionophore or due to low Ply production inside PCVs not only triggers LC3 lipidation without assistance from other autophagic components, but also prolonged the persistence of SPN in the host cells.

**Fig 5:**
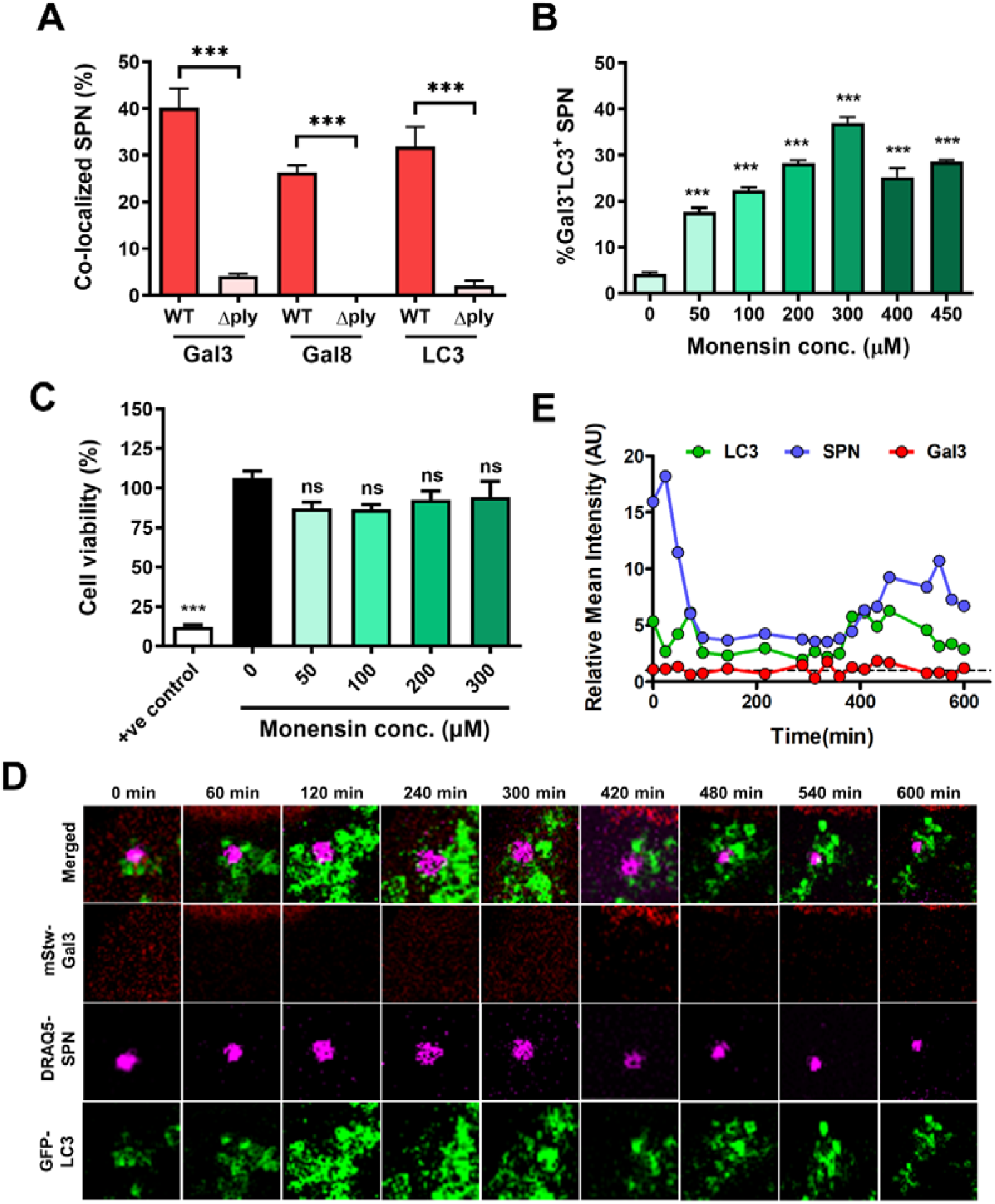
Induction of osmotic imbalance triggers decoration of Δ*ply* mutant with LC3 and improves intracellular persistence. **(A)** Percentage of association of Δ*ply* mutant with Gal3, Gal8 and LC3 in comparison to WT SPN. Statistical analysis was performed using Students t-test. \*\*\**p*<0.001. **(B)** Quantitative analysis showing increased association of Δ*ply* mutant with LC3 in absence of damage sensing marker Gal3, upon treatment with different conc. of monensin. n ≥ 100 bacteria per coverslip. Data are presented as mean ± SD of triplicate experiments. Statistical analysis was performed using one-way ANOVA followed by Tukey’s Multiple Comparison Test. \*\*\**p*<0.001. **(C)** Cell viability at different conc. of monensin. Triton X-100 (0.1%) served as a positive control. Statistical analysis was performed using one-way ANOVA followed by Tukey’s Multiple Comparison Test. ns, non-significant, \*\*\**p*<0.001. **(D)** Representative time-lapse montage of Δ*ply* mutant showing prolonged persistence post monensin treatment (50 μM). A549s stably expressing mStrawberry-Gal3 and GFP-LC3 were infected with DRAQ5 stained Δ*ply* mutant. Following monensin treatment cells were monitored using confocal microscope at an interval of 60 min for extended hours. The stills correspond to the movie shown as S6 Movie. **(E)** Temporal quantification of Gal3, LC3 and SPN fluorescence intensities either relative to fluorescence in the cytosol (*I_PCV_/I_Cyt_*) (for LC3 and Gal3) or fluorescence signal of A549 nuclei (*I_SPN_/I_Nuc_*) (for SPN). *I_PCV_/I_Cyt_* values were always found to be close to 1.0 for Gal3, exhibiting no association of Gal3 with PCVs. Contrarily *I_PCV_/I_Cyt_* for LC3 and *I_SPN_/I_Nuc_* for SPN were > 1.0, revealing prolonged persistence of SPN inside LC3 marked vacuole for extended hours.

### Mathematical modelling of Ply expression triggering varied outcomes

Though, we have been able to clearly delineate the role of Ply in governing the fate of intracellular SPN, the quantitative details of *ply* gene expression pattern triggering such outcomes remained unknown. To build a simple mathematical model depicting relation of SPN lifespans with *ply* expression, we hypothesize that pore formation on PCVs and subsequent escape of SPN to cytosol is a result of Ply accumulation crossing certain threshold. Thus, apart from the process of protein production, all other associated biophysical steps (such as details of deposition on endosomal membrane, oligomerization etc.) are ignored at the first approximation. Such reduced picture of an effective first passage process of protein threshold crossing has been earlier reported to understand the timings of many significant cellular events like lysis, cell division, and kinetochore capture [16–23].

When Ply accumulation in the proximity of the PCV wall causes damages on it, Gal8 molecules are recruited from the cytosol on those damaged points in random times *t_g8_*. The histogram for *t_g8_* for the vacuolar and cytosolic cases, respectively, (from Fig. 1D and Fig. 2D), are plotted in Suppl. Fig. 4A and 4B, respectively. They follow exponential distributions with decay constants ~ 0.0124 min^−1^ and 0.0073 min^−1^ (consistent with the mean times ~80 min and ~137 min as shown in Fig. 1D and 2D). The distributions are expected to be exponential as the Gal8 attachment to a PCV is likely to be a single-step process.

For the SPN population subset which die in the cytosol after escape through the pores in PCV, histograms for *t_C’_* (total time of residence of the SPN inside PCV before escape to cytosol) and *t_p_* (time of larger pore formation after Gal8 attachment enabling escape to cytosol) are shown in Figs. 6A and 6B, respectively (using data from Fig. 2D-E). We assume that during the time periods *t_C’_* or *t_p_*, the Ply number increases from some initial value *N_0_* to some final value *N* due to Ply translation. The number (*N-N_0_*) of Ply is expected to be larger for *t_C’_* compared to *t_p_*. This number (*N-N_0_*), the protein production rate *k*, and the mean burst size *b*, are unknown parameters which can be obtained by fitting the experimental histogram to the theoretical formula in Eq. (1) (see materials and methods). By best-fitting (see materials and method) the experimental data for *t_p_* (Fig. 6B), we estimated the parameter values of Ply burst size (*b*) to be 2, the rate of Ply production (*k*) to be 0.107 min^−1^ and the number of Ply required to reach the threshold (*N-N_0_*) to be 13. We see that the theoretical formula (in solids red line) fits the experimental data quite well and reproduces the skewed non-Gaussian nature to the right of the maximum. With our fitted values of *b, k*, and (*N-N_0_*), Eq. (3) (in materials and methods) we get the theoretical standard deviation σ(*t_p_*) = 38.7 min (close to the experimental σ ~ 44.7 min). When we fit the experimental data for *t_C’_* (Fig. 6A) in a similar way with the theory, the value (*N-N_0_*) turns out to be 35. This provides us with a rough estimate of intravacuolar number of Ply (35-13 = 22) accumulated during the time (~ *<t_g8_>* = 137 min) of Gal8 attachment. Thus, such a model not only provided the rate of Ply synthesis (*k* ~ 0.107 min^−1^), but also enabled quantitative estimates of Ply accumulated during Gal8 attachment (~ 22) and larger pore formation on PCV membrane post Gal8 attachment to facilitate cytosolic escape (~13).

**Fig 6:**
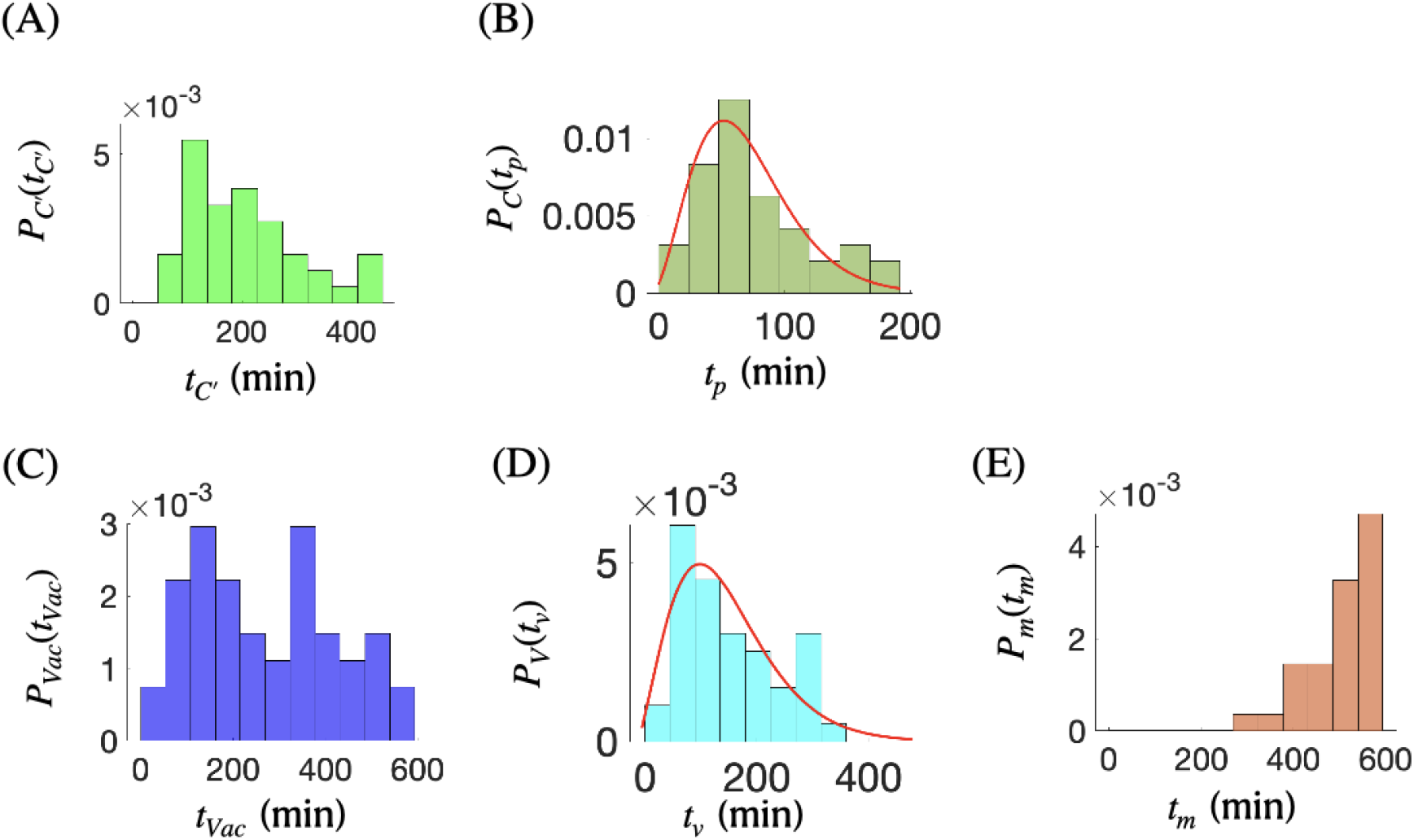
Fitting of experimental histograms gives the quantitative estimates of Ply growth rate (*k*), Ply burst size (*b*) and the threshold Ply number (*N-N_0_*) required for accomplishing an event in the degradation pathway. **(A)** Normalized histogram of total time of pore formation, *t_C’_* (in min), on PCV for escape to the cytosol, by high-Ply producing subpopulation among WT SPN. **(B)** Normalized histogram of cytosolic escape time *t_p_* (in min), after Gal8 recruitment. The continuous red line represents the theoretical best-fitted curve (*b* = 2, (*N-N_0_*) =1 3, *k* = 0.107 min^−1^) to the experimental histogram. **(C)** Normalized histogram represents the total degradation time *t_Vac_* (in min) of WT SPN subpopulations with moderate or low Ply production within PCV. **(D)** Normalized histogram of degradation time *t_v_* (in min) of WT SPN within vacuoles is plotted for times < 400 min. The theoretically best-fitted line in red (*b* = 2, (*N-N_0_*) = 10, *k* = 0.0412 min^−1^) is shown against the cyan histogram. **(E)** Normalized histogram of degradation times *t_m_* (in min) measured after the LC3 recruitment, for LC3^+^Gal3^-^ marked subpopulation among the SPN:Ply-Low strains.

The histogram (using data from Fig 2F) of timescales of SPN degradation in the cytosol after escape from PCVs (*t_c_*), are shown in Suppl. Fig. 5, but since pneumococcal degradation post cytosolic escape are independent of Ply, we do not attempt to model the statistics of these timescales.

In the case of vacuolar degradation, the bacteria eventually degrade within the Gal8 marked autophagosomes. We hypothesize that due to relatively low Ply production, this subset of SPN cannot create significantly large pores on PCV membrane to escape to cytosol. So we expect that our theoretical model would provide threshold number of Ply in this case to be lower compared to the cytosolic SPN population discussed above. We first plot the histograms of the total vacuolar degradation times (*t_Vac_ = t_g8 +_ t_v_*) in Fig. 6C (using Fig. 1F), as well as the degradation times *t_v_* after Gal8 attachment in Fig. 6D (using Fig. 1E). The histogram in Fig. 6C exhibits bi-modality, suggesting existence of two separate peaks at different time points. This could presumably be due to distinct subpopulations of WT SPN which produces Ply at distinct rates within different vacuoles, and because of which they have shorter or longer lifetimes. This is supported by our experimental study of a low-Ply expressing mutant (SPN:Ply-Low) having distinct markers Gal3^+^LC3^+^, Gal3^+^LC3^-^ and Gal3^-^ LC3^+^ associated with different SPN subpopulations. Among these, we found that Gal3^-^LC3^+^ marked subset is the majority and longest living with lifetimes (*t_m_*) ranging from ~250 min to ~600+ min with a significant rise beyond 400 min (Fig. 6E). Hence, in Fig. 6D we intentionally plot histogram of time scales below 400 min, which presumably represent moderate-Ply producing subpopulation. Those above 400 min (corresponding to extremely low-Ply producing subsets) have been ignored. Fitting the experimental histogram (Fig. 6D) with the theoretical formula Eq. (1) [materials and method], we obtained the mean Ply burst size (*b*) to be 2, the rate of Ply production (*k*) to be 0.0412 min^−1^ and increment of Ply number (*N-N_0_*) = 10, during its lifespan following Gal8 attachment and its eventual death. The theoretical fit is shown in solid red line in Fig. 6D. With the fitted values of *b, k*, and (*N-N_0_*), we get (using Eq.3 in materials and method) a standard deviation of 89.2 min (compared to the experimental σ ~ 85.5 min).

Just like cytosolic population, a fit to the experimental data for *t_Vac_* (Fig. 6C) yields the value (*N-N_0_*) = 19. Again, this helps in finding a rough estimate of number of Ply (19-10 = 9) accumulated in the vacuole before (~ *<t_g8_>* = 80 min) of Gal8 attachment. Critically, our analysis suggests: (i) the rate of Ply synthesis in case of SPN subset undergoing vacuolar degradation is 2.6 fold lower (0.107 min^−1^ vs 0.0412 min^−1^) compared to SPN subpopulation undergoing cytosolic death, and (ii) the number (~19) of Ply accumulated in case of SPN subset undergoing vacuolar degradation is almost half of the number of Ply (~35) needed for the cytosolic escape.

## Discussion

Though pneumococci is generally regarded as an extracellular pathogen, in recent years it has been evident that SPN occasionally establish intracellular niches within the body to evade immune surveillance and disseminate within the host [6, 38]. Particularly pertinent to this are observations of its intracellular replication inside splenic macrophages and cardiomyocytes, as well as prolonged persistence in the lower respiratory tract [6, 8]. Central to its intracellular life is expression of pore forming toxin pneumolysin, which has been shown to form variable-sized pores on biological membranes. Though Ply’s differential pore-forming ability is directly dependent on the monomer concentration of the toxin in *in vitro* conditions [39–42], reports depicting correlation between its expression levels and pore dynamics, fostering its intracellular life, is not available.

Few other pore-forming proteins, such as Holin, are reported perforate cell membrane only after accumulating to a threshold level [17]. Such proteins are usually uniformly distributed in a relatively mobile state in the cell membrane. A sudden transition from this uniformly distributed state to accumulation at specific sites occurs during membrane damage which eventually results in formation of holes in the membrane. Therefore, to calculate the membrane perforation rate of a protein toxin, it is necessary to keep track of the timescale of such transition [43]. On similar lines, we hypothesized that various events observed by us, ranging from vacuolar degradation to cytosolic escape and subsequent cytosolic degradation, have happened due to Ply accumulation inside PCVs reaching a threshold value. Firstly, we quantitatively studied lifespans of SPN subpopulations that had these distinct fates. Both high and moderate Ply producing subpopulations were marked with Gal8--the corresponding timescales for Gal8 recruitment were ~137 and 80 min, respectively. But post Gal8 attachment, while the vacuolar subset stayed alive for a further ~104 min, the High-Ply variants escaped to the cytosol in relatively shorter time (~70 min). On the contrary, the very low Ply producing variants showed extremely prolonged persistence (>400 min). Our main aim in this manuscript was to correlate the shorter vs longer lifespans of distinct SPN subpopulations with the corresponding amount of Ply accumulation inside the vacuoles. Towards this end we quantified important timescales for two events, the cytosolic escape and the vacuolar degradation events, associated with the degradation pathways inside the host cell.

By comparing these events with a simple mathematical gene expression model, where protein numbers reaching a threshold value could trigger a biochemical function, we evaluated the rate of protein production, the protein burst size and threshold number, from experimental histograms. Our findings suggested that the Ply accumulation is proportional to the protein production rate (*k_c_* and *k_v_* for cytosolic and vacuolar life-stages, respectively). Critically, we found that *k_v_* is 2.6 times slower than *k_c_*, strengthening our speculation. At the same time, the Ply threshold numbers associated with the cytosol escaped population is ~2 fold higher than in the case of SPN subset degrading within the vacuole. It could be speculated that due to slow production rate of Ply in case of vacuolar SPN population, the time taken for threshold Ply synthesis could be sufficiently long, providing the host a window of opportunity to sequester the Gal8 marked damaged endosomes in autophagosomes. While for the cytosolic population, the high rate of Ply synthesis could quickly lead to catastrophic damage of Gal8 marked endosomes, triggering SPN’s escape to cytosol even before the host could sequester it in autophagosomes. We would like to highlight that such quantitative estimates of intravcuolar toxin accumulation triggering engagement with diverse host intracellular defense mechanisms promoting distinct phenotypic outcomes is a novel finding.

Our results also suggested that Gal3 can detect smaller pores and is possibly recruited to the damaged endosomes first, compared to Gal8. Unlike other galectins, not only Gal3 is widely distributed in different tissues (Gal8 is also widely distributed), it is the only galectin that has a long unique N-terminal domain attached to a single carbohydrate recognition domain (CRD) [44]. Gal3 can exist as monomer or can associate via the non-lectin domain into multivalent complexes up to a pentameric form [45]. It is speculated that this allows Gal3 to bridge effectively between different ligands and form adhesive networks. Though diverse role of Gal3 in mediating endomembrane repair or targeting the damaged endosomes towards autophagy is already known [36], how Gal3 switches between these roles remain elusive. On one hand Gal3 is known to interact with ESCRT complex including Alix to augment endomembrane repair, on the other it interacts with TRIM16, an autophagy receptor, for efficient sequestration of damaged endosomes or lysosomes [46]. We also similarly observed that subsets of the low Ply producing SPN population, beyond Gal3 decoration either associate with autophagy marker LC3 or remain free of LC3 (repair pathway). However, irrespective of the routes adopted, both pathways finally resulted in bacterial degradation, possibly following fusion with lysosomes.

On the contrary, the unique SPN population subset which exhibited prolonged intracellular persistence, remain devoid of Gal3 as well as other damage sensing markers throughout its lifetime, but are unusually decorated with autophagy marker LC3. Recently, infection by few pathogens, such as *L. monocytogenes, A. fumigatus* etc. are reported to induce decoration of pathogen containing phagosomes by LC3 in a process called LC3-associated phagocytosis (LAP) [47–49]. Such processes have been shown to result in enhanced degradation of the cargo as well as antigen presentation by MHC-II molecules [50]. In contrary to the classical autophagy process where LC3 is conjugated to phosphatidylethanolamine (PE), in LAP or related non-canonical autophagy pathways LC3 lipidation is observed to happen on phosphatidylserine (PS) [51]. It is assumed that such conjugation may provide a “molecular signature” for non-canonical autophagy, enabling its distinction from closely related, parallel pathways. However, the LC3^+^Gal3^-^ PCVs we observed are not generated by LAP as recruitment of LC3 to these endosomes occurs even after inhibition of PI3K, a prerequisite for induction of LAP (data not shown). Few pathogens possessing pore forming toxins are also known to trigger a variant of non-canonical autophagy which is distinct from LAP [52, 53]. This process, termed as pore-forming toxin-induced non-canonical autophagy (PINCA), can also be initiated by the needle-like Type three secretion system (T3SS) of *S. flexneri* or *S. typhimurium* [54]. In these cases, ion or osmotic imbalance induced by bacterial toxins or T3SS is responsible for LC3 lipidation of pathogen containing vacuoles. This is typified by *H. pylori* toxin, VacA which induces osmotic imbalance to cause such unconventional LC3 lipidation on intact phagosomes [37]. Drawing parallel with this, we speculated that the long persisting PCVs have originated due to extremely low Ply expression which resulted in very minute damage to the endomembrane. Such disruptions are not capable of recruiting any damage sensors, but are adept in inducing pH or ionic imbalance. Indeed, we could trigger decoration of Δ*ply* mutant containing PCVs with LC3 following treatment with an ionophore, proving the key role of ion or osmotic imbalance induction in marking of perturbed endomembrane with LC3.

However, the LC3^+^Gal3^-^ PCVs we observed, differ fundamentally with other pathogen containing vacuoles that are marked by non-canonical autophagy in persistence time and ultimate fate. For example, in PINCA, LC3-positive phagosomes fused more often with lysosomes indicating that PINCA promotes phagolysosomal fusion [55]. It is also speculated that PINCA might represent an attempt of macrophages to repair damaged phagosomal membranes as last resort against pathogens, which are yet to escape from the phagosome [56]. Contrary to this, in our case tiny pores produced by low Ply expression, help SPN to sustain for prolonged periods. This could be correlated to a situation with *L. monocytogenes* where lesser listeriolysin O (LLO) production gave rise to a unique subpopulation that resides inside the phagosomes and persists by lowering their metabolic activity. Such <u>S</u>pacious <u>L</u>isteria <u>C</u>ontaining <u>P</u>hagosomes (SLAPs) serve as a replicative niche for *Listeria* by maintaining a neutral intraluminal pH [3]. This highlights that residing inside a vacuole by lowering metabolic activity can be a pathogenic counter-strategy to evade host defences. Our findings also point towards a similar situation where SPN deliberately lowers its toxin production to ensure longer persistence without triggering inflammation. Prolonged persistence associated with neutralization of bacteria-bearing lysosomes is reported to trigger lysosome exocytosis, resulting in expulsion of exosome-encased bacteria [57]. Though such a process is touted to be a cell autonomous defence program to clear recalcitrant pathogens, these could potentially be exploited by pathogens for egressing out of the hostile intracellular environment. Whether pneumococci follow similar suit for egressing out of lung epithelium, facilitating onward transmission, remains to be explored.

## Acknowledgement

We acknowledge the biosafety level 2 facility and confocal microscopy facility at IIT Bombay. S.S. acknowledges financial support from IIT Bombay. D.D. acknowledges financial support from Science and Engineering Research Board (SERB) India (Grant No. MTR/2019/000341). A.B. acknowledges research funding from Science and Engineering Research Board (SERB), Government of India (Grant no. SPR/2019/000808). The funder had no role in study design, data collection and interpretation, or the decision to submit the work for publication.

## Notes

### Competing Interest Statement

The authors have declared no competing interest.

