## Supplemental Material for "Quantification of differential toxin expressions and their relation to distinct lifespans of bacterial subpopulations associated with diverse host immune mechanisms"

**This file contains:**

Supplementary Figures 1 to 5

Supplementary Movies 1 to 6

**Supplementary Fig. 1.**

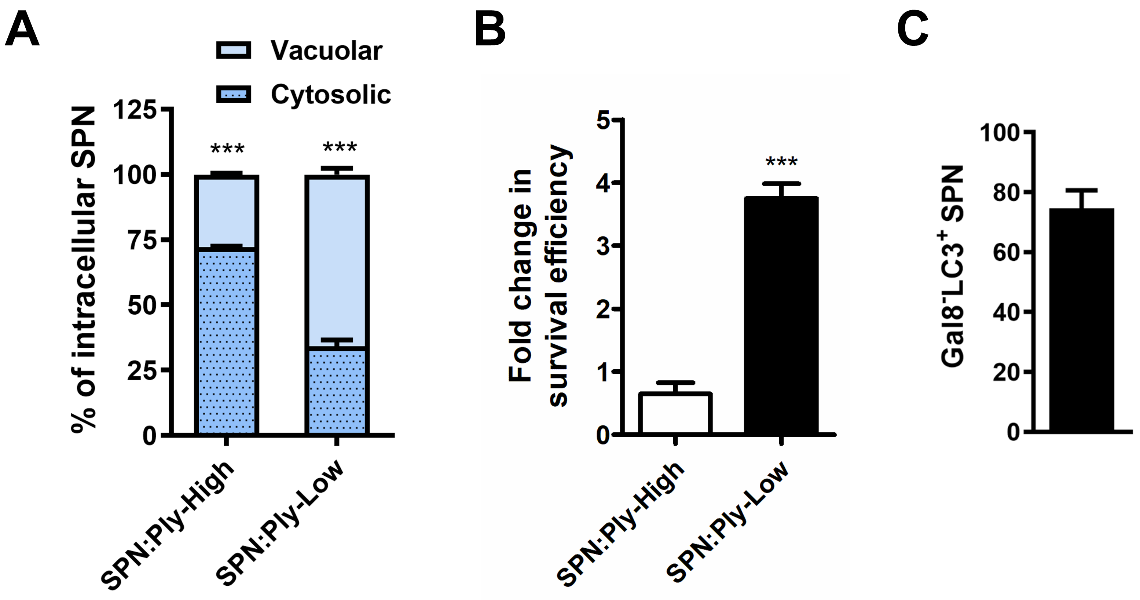

**Intracellular fate of SPN is governed by differential Ply expression.**

**(A)** Intracellular location of WT:Ply-High and WT:Ply-Low SPN strains at 8 h post infection in A549 cells. Flip-TR positive PCVs are considered as vacuolar SPN population, while Flip-TR^–^ are considered to be cytosolic. n > 100 bacteria per coverslip. Data are presented as mean ± SD of triplicate experiments. **(B)** Fold change in intracellular survival efficiency of SPN strains expressing low (SPN:Ply-Low) or high (SPN:Ply-High) levels of Ply at 8 h post infection in A549 cells. **(C)** Percentage of SPN:Ply-Low strain containing vacuoles marked with LC3 but devoid of Gal8 (LC3^+^ Gal8^-^) at 8 h post infection. Statistical analysis was performed using two-way ANOVA followed by Bonferroni test (**A**) or students t-test (**B**). ***p<0.001.

**Supplementary Fig.** **2.**

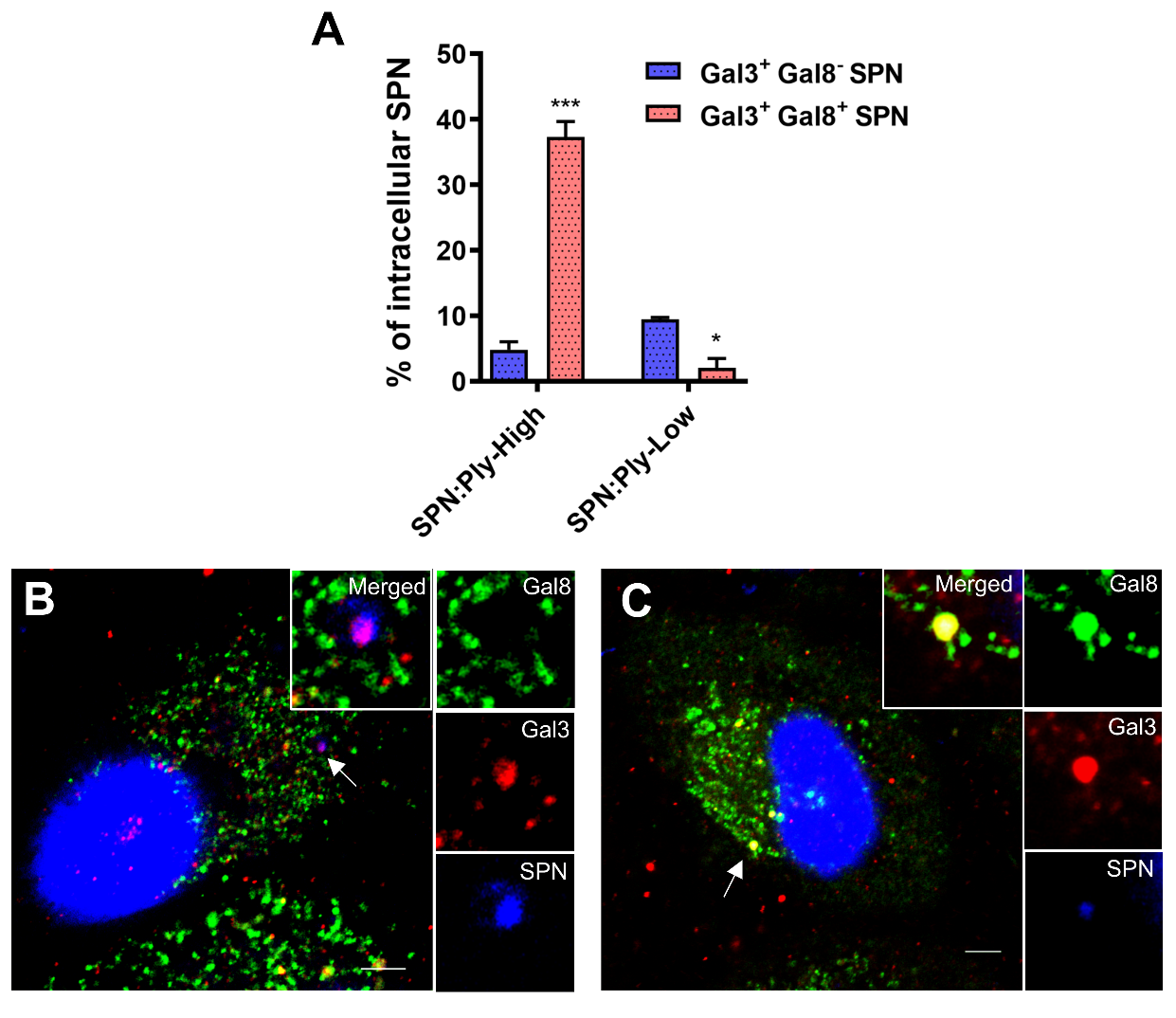

**Low Ply expressing SPN has distinct subpopulations.**

**(A)** Quantification of association of endomembrane damage sensing markers Gal3 and Gal8 with SPN:Ply-High and SPN:Ply-Low strains at 8 h p.i. n > 100 bacteria per coverslip. Data are presented as mean ± SD of triplicate experiments. Statistical analysis was performed using two-way ANOVA followed by Bonferroni Test. *p<0.05; ***p<0.001. **(B-C)** Confocal images of Gal3^+^ Gal8^-^ SPN **(B)** and Gal3^+^ Gal8^+^ SPN **(C)**. A549s stably expressing YFP-Gal8 were infected with WT SPN (blue) and stained with anti-Gal3 Ab. Arrowhead designates bacteria shown in insets. Scale bar, 5 μm.

**Supplementary Fig. 3.**

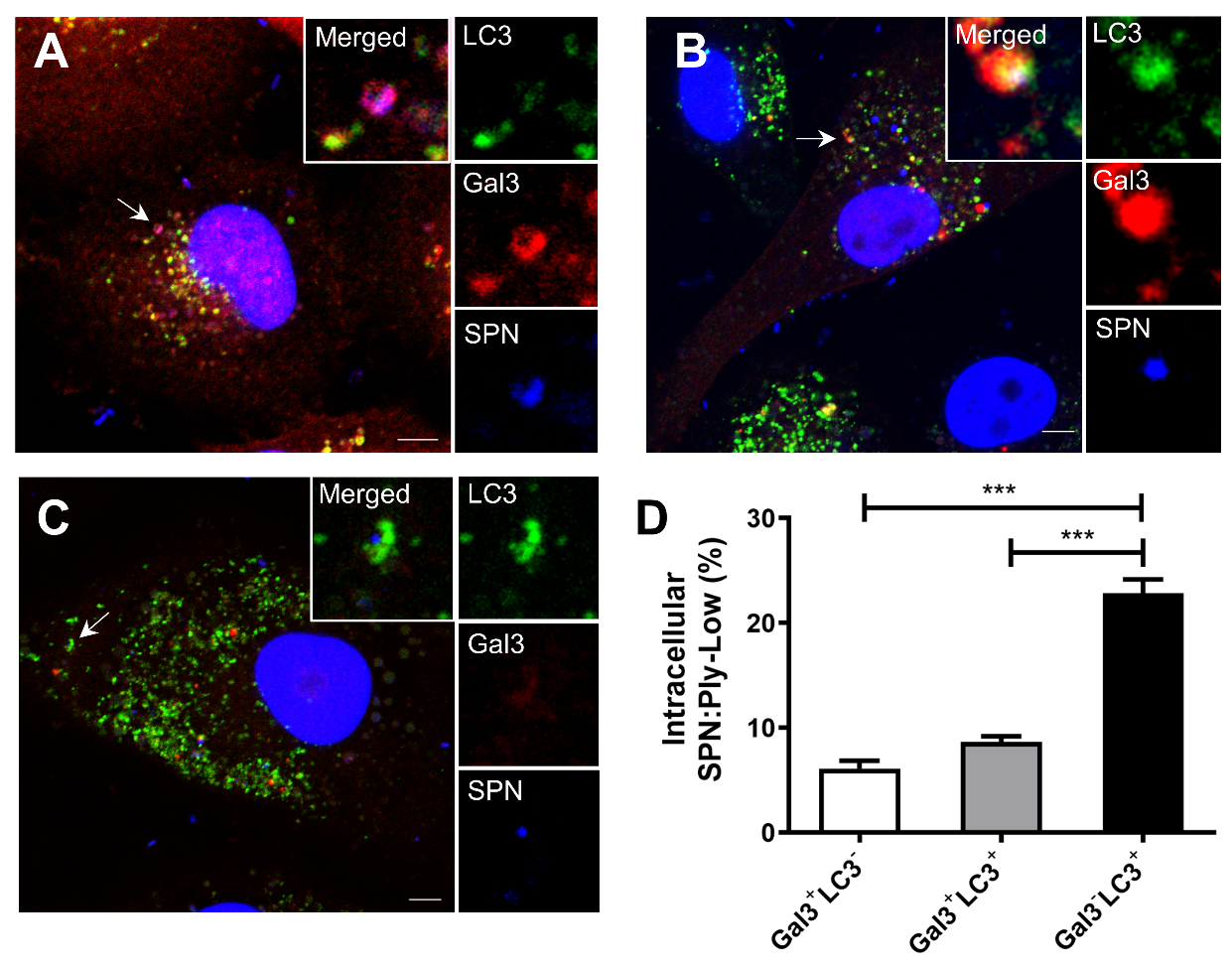

**Low Ply expressing SPN has distinct subpopulations.**

**(A-C)** Confocal images of different population of SPN:Ply-Low strain inside A549s at 8 h p.i. **(A)** Gal3^+^ LC3^-^, **(B)** Gal3^+^ LC3^+^, **(C)** Gal3^-^ LC3^+^. A549s stably expressing mStrawberry-Gal3 and GFP-LC3 were infected with WT SPN (blue). Arrowhead designates bacteria shown in insets. Scale bar, 5 μm. **(D)** Quantification of different population subsets of intracellular SPN:Ply-Low strain at 8 h p.i. for association with Gal3 and LC3. n > 100 bacteria per coverslip. Data are presented as mean ± SD of triplicate experiments. Statistical analysis was performed using one-way ANOVA followed by Tukey's Multiple Comparison Test. ***p<0.001.

**Supplementary Fig. 4.**

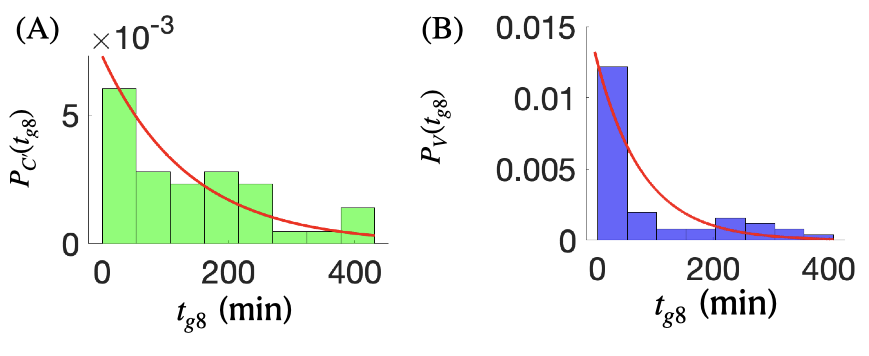

**Fitting of experimental histograms of Gal8 recruitment times give exponential distribution representing a single-step process.**

Normalized histogram of Gal8 recruitment times *t_g8_* (in min) on the PCVs, containing WT SPN, for the cytosolic (**A**) and vacuolar (**B**) degradation pathways, respectively. The solid red lines represent the exponential curves with decay constants 0.0073 min^-1^ and 0.0124 min^-1^ in (**A**) and (**B**), respectively.

**Supplementary Fig. 5.**

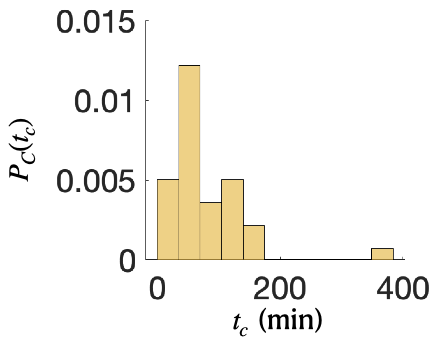

**Histogram of cytosolic residence time of SPN following vacuolar escape.**

Experimental histogram (n = 40) of cytosolic residence time *t_c_* of SPN after escape from autophagosome obtained from Fig. 2F.

**S1 Movie: Vacuolar killing of SPN.** Time lapse fluorescence imaging demonstrating degradation of tRFP expressing SPN (red) following infection in A549s stably expressing YFP-Gal8 (green). Live cell imaging was performed under spinning disc confocal microscope starting at 60 min p.i. The video depicts degradation of SPN while being associated with Gal8 signal. Time interval, 12 min; scale bar, 1.5 µm.

**S2 Movie: SPN’s cytosolic escape and eventual death.** Time lapse imaging of tRFP expressing SPN (red) infected A549s stably expressing YFP-Gal8 (green). Live cell imaging was performed under spinning disc confocal microscope starting at 60 min p.i. The video depicts degradation of SPN in cytosol after it escapes from Gal8 marked endosomes. Time interval, 12 min; scale bar, 1.5 µm.

**S3 Movie: Galectin 3 mediated intravacuolar non-autophagic killing of SPN.** Live fluorescence imaging of DRAQ5 stained SPN’s (magenta) degradation of inside A549s stably expressing mStrawberry-Gal3 (red) and LC3-GFP (green). Live cell imaging was performed under laser scanning confocal microscope starting at 30 min p.i. The video depicts killing of SPN while being in damaged vacuole marked with Gal3, but devoid of LC3. Time interval, 20 min; scale bar, 1.5 μm.

**S4 Movie: Galectin 3 driven autophagic killing of SPN.** Live fluorescence imaging of A549 cells stably expressing mStrawberry-Gal3 (red) and LC3-GFP (green) following infection with DRAQ5 stained SPN’s (magenta). Time lapse imaging was performed under laser scanning confocal microscope starting at 30 min p.i. The video depicts SPN degradation in autophagosomes following endomembrane damage mediated association of Gal3. Time interval, 25 min; scale bar, 1.5 μm.

**S5 Movie: Persistence of SPN in non-canonical LC3 marked compartments.** Live fluorescence imaging of A549 cells stably expressing mStrawberry-Gal3 (red) and LC3-GFP (green) following infection with DRAQ5 stained SPN’s (magenta). Time lapse imaging was performed under laser scanning confocal microscope starting at 30 min p.i. The video depicts association of SPN with LC3 without being marked with Gal3. It also portrays prolonged persistence of SPN in these only LC3 marked non-canonical autophagosomes. Time interval, 65 min; scale bar, 1.5 μm.

**S6 Movie: Induction of ionic imbalance triggers LC3 association and prolongs SPN’s intracellular survival.** Time lapse fluorescence imaging of A549 cells stably expressing mStrawberry-Gal3 (red) and LC3-GFP (green) following infection with DRAQ5 stained Δ*ply* mutant SPN strain (magenta). Live imaging was performed under laser scanning confocal microscope starting at 30 min p.i. The video depicts association of Δ*ply* mutant SPN with LC3 without being associated with Gal3 following treatment with an ionophore, monensin. It also portrays prolonged persistence of SPN in these only LC3 marked non-canonical autophagosomes. Time interval, 60 min; scale bar, 1.5 μm.
